# Targeting breast cancer senescence in 3D models of bone metastasis

**DOI:** 10.1101/2025.06.26.661856

**Authors:** Eleane C.B. Hamburger, Mansoureh Mohseni Garakani, Saleh Alfaisali, Jean A. Ouellet, Michael H. Weber, Livia Garzia, Lisbet Haglund, Derek H. Rosenzweig

## Abstract

Chemotherapeutic treatment of breast cancer with Doxorubicin can induce tumor and stromal cell senescence leading to therapy-resistance. Senescence-associated secretory phenotype (SASP) promotes secretion of pro-inflammatory and tumorigenic factors causing systemic inflammation. Combined, this can result in immune suppression, tumor growth and secondary spread of cancer. Targeting and removing senescent and cancerous cells using a combination of chemotherapeutic and senolytic drugs may reduce systemic inflammation, improve therapeutic efficacy, and prevent metastasis. Treatment of both triple-negative breast cancer (MDA-MB-231) cells, and primary spine osteoblasts 0.25 µM Doxorubicin showed significant induction of senescence indicated by p21 positive cells. Doxorubicin and senolytics (RG-7112, o-Vanillin) treatment of mono-culture and co-culture spheroids showed a significant additive effect on decreased tumor sphere viability and growth. This was correlated with decreased p21 and Ki67 proliferation marker in both the breast cancer and osteoblast cells. In all cases, combined Doxorubicin and senolytics significantly reduced sphere size and cancer cell outgrowth, indicating reduced metastatic potential. Future chemotherapeutic treatment of breast cancer patients may be optimized by adding senolytic drugs to more effectively clear tumors and help regenerate surrounding stroma tissue such as in the bone metastatic environment.

## Introduction

Metastatic tumors of the spine are the most common type of bone metastasis. These tumors often originate from primary breast, prostate, lung, and kidney cancers [1-5]. With advancements in medical, radiation and surgical oncology, these patients have a longer life-expectancy post-diagnosis [6], and the societal disease burden is growing [7, 8]. Indeed, breast to spine metastasis is one of the most common metastases with 85% incidence; it is also among the leading causes of death in women with metastatic breast cancer [9]. Metastatic spine disease is often accompanied by neurological and structural deficits leading to poor health related quality of life driven by pain, instability or paralysis [10, 11]. This has been mostly attributed to osteolysis within the spine bones, where the receptor activator of nuclear factor κB (RANK), its ligand (RNAKL) and osteoprotegerin (OPG – inhibitor of RANKL) signaling becomes imbalanced toward osteoclasts activity [12]. Current treatments for metastatic spine lesions include acute radiotherapy and when advisable, surgery. Patients with advanced disease are often also given systemic chemotherapy which can increase systemic inflammation and disrupt immune function[13]. Considering these patients were also given standard systemic chemotherapy during their primary tumor care, the new tumor microenvironment may become enriched with senescent cells further promoting tumor progression and drug resistance.

Both radiotherapy and chemotherapy are known for inducing high levels of stress induced cellular senescence [14]. Cellular senescence, or irreversible cell cycle arrest, is characterized by changes in gene expression, metabolism, and anti-apoptotic pathways[15]. Doxorubicin is anthracycline which blocks DNA topoisomerase 2 and is a standard chemotherapeutic for treating advanced breast cancer. Doxorubicin is toxic to normal tissues and organs, and clinicians are forced to use suboptimal doses to minimize side effects[16, 17]. This can result in incomplete removal of cancer cells, as well as therapy resistance and increased cellular senescence in both cancer and normal cells in the tumor microenvironment and beyond[18]. Although senescence may initially halt cancer cell proliferation, the adoption of a senescence-associated secretory phenotype (SASP), results in the secretion of pro-inflammatory and pro-tumorigenic factors. In later stages, senescence can paradoxically drive the growth and spread of the cancer possibly through SASP factors contributing to chronic, systemic inflammation, immune suppression as well as chemoresistance[19]. Moreover, the senescence of surrounding stroma gives negative feedback on the tumor microenvironment and vice-versa[20]. Therefore, removing or suppressing both senescent cancer and stromal cells may be useful in targeting the metastatic breast-to-bone niche and improving chemotherapeutic efficacy for these patients.

Two classes of senotherapeutic (senescence-targeting) drugs have been developed called senolytics and senomorphics. Senolytic drugs induce apoptosis of the senescent cells, while senomorphics suppress the pathological phenotypic trait of the senescent cell, such as secretion inflammatory and catabolic factors[21]. Our group and others have shown that senolytics such as RG-7112, and senolytics and senomorphics such as o-Vanillin, are capable of suppressing musculoskeletal pathologies and improving immune function and tissue homeostasis[19, 22-25]. RG-7112 is a member of the p53/MDM2 complex inhibitors. It is FDA approved for acute myeloid leukemia and previously underwent clinical trials for the treatment of various cancers[26]. High doses required for anti-cancer activity unfortunately lacked efficiency and showed hematological toxicity[27, 28]. O-Vanillin is a natural compound whose metabolites have both anti-inflammatory and senolytic effects[29]. Interestingly, o-Vanillin has been linked to anti-metastatic, anti-proliferative and anti-angiogenic properties in cancer related studies[30]. Whether these two compounds can effectively remove senescent cells and their related SASP to improve efficacy of chemotherapy remains unclear.

In this study, we set out to assess whether combined senotherapy and chemotherapy could overcome the negative impact of therapy-induced senescence in a breast-to-bone microenvironment model. We hypothesized that combining RG-7112 and o-Vanillin will remove therapy induced senescent cells from the stromal and tumor compartments, allowing Doxorubicin to more effectively block tumor sphere growth, viability, and migration. Understanding the additive or synergistic effects of this combined therapeutic approach may help in combating metastatic disease in the spine and provide avenues toward personalized patient care for this cancer and potentially others.

## 2. Materials and Methods

### 2.1. Cell isolation and *in vitro* culture

#### 2.1.1. Tissue collection and bone cells isolation

Primary spine osteoblasts were isolated from human lumbar spine tissue with informed consent in collaboration with Transplant Quebec and approved by the ethical review board at McGill University (Institutional Review Board, IRB #2019-4896). The cortical bone chips cut from vertebral bodies were approximately 3-5mm^3^ and washed with 2X phosphate-buffered saline solution (PBS, Sigma-Aldrich, Oakville, ON, Canada) supplemented with Primocin (InvivoGen, San Diego, CA, USA) and Fungizone (Sigma-Aldrich, Oakville, ON, Canada) as previously described[31]. The bone chips isolated are then placed on a moving rocker at 37°C in 1.5mg/mL collagenase type II solution (Gibco, Canada) in basal media for 12-14 hours to remove any remaining soft tissue. Bone chips were re-washed in the 2X PBS supplemented with antibiotics and plated in T75 flasks with Dulbecco’s Modified Eagle Medium 4.5g glucose (DMEM) (Gibco, Canada, #12430062) media supplemented with 10% heat inactivated fetal bovine serum (FBS (Gibco, Canada, #12483020)) and 1% penicillin/streptomycin (PS) (Gibco, Canada, #15140122)) and incubated at 5% CO_2_ 37°C to allow for osteoblast cells to crawl out of the bone chips. Cells were passaged at 80% confluency and used from passage 1-4. Tissue demographics can be seen in Table 1.

**Table 1.** Characterization of vertebral tissue donor demographics.

| Age | Sex | Cause of Death |
| --- | --- | --- |
| 18 | F | Drowning |
| 33 | F | Trauma |
| 54 | F | Cardiac Arrest |
| 60 | F | Cerebral Hemorrhage |
| 73 | F | Stroke |
| 16 | M | Overdose |
| 19 | M | Overdose |
| 20 | M | Trauma |
| 31 | M | Cerebrovascular Accident |
| 65 | M | Cerebral Hemorrhage |

#### 2.1.2. Cell culture and seeding

Primary spine osteoblasts procured as mentioned above and green fluorescent-protein (GFP) expressing epithelial breast adenocarcinoma cell line GFP-MDA-MB-231 provided by the laboratories of professor M. Park at McGill University, were cultured in T75 flasks (Starstedt, TC Flast T75, Stand, Vent. Cap, Germany) in basal media, high-glucose DMEM (Gibco, Canada) supplemented with 10% FBS (Gibco, Canada) and 1% PS (Gibco, Canada). Cells were placed in incubators 37°C and 5% CO_2_ until 80% confluency was reached for passaging.

a. Standard 2D culture Chamber slides (Nunc™ Lab-Tek II™ ThermoScientific) were incubated with DMEM full serum media for 24 hours then seeded with 20,000 MDA-MB-231 cells per well or 15,000/well primary spine osteoblasts and allowed to adhere overnight at 37°C, 5% CO_2_.
b. Spheroid 3D culture/co-culture For spheroid generation, twenty-thousand MDA-MB-231 cells suspended in 100ul DMEM media (10% FBS, 1%PS) were seeded per well of a Nunc™ 96-well, Nunclon Delta-Treated, U-Shaped-Bottom Miroplate (ThermoFisher Scientific, Toronto, ON, Canada). Spheroid generation protocol is as based on MDA-MB-231 Cell Line Spheroid Generation and Characterization for HT Assays (ThermoFisher Scientific, Toronto, ON, Canada). For co-culture experiments, following 24 hours of spheroid generation, osteoblasts are seeded at 3 million/mL of supplemented DMEM media mixed with PureCol® EZ Gel Bovine Collagen Solution Type I Collagen 5mg/mL (CELLINK, BICO, Göteborg, Sweden) for a 50/50 dilution. Cells mixed in diluted collagen are seeded at the density of 300,000 cells per well and re-centrifuged based on MDA-MB-231 Cell Line Spheroid Generation and Characterization protocol for HT Assays (ThermoFisher Scientific). The plate is placed in an incubator to allow collagen to be crosslinked at 37°C for 30 minutes before 100ul media is overlayed atop and incubated for 3 days to mature. On day 3 spheroids were transferred into a 48 non-adherent well plate (Geiner CELLSTAR®, multi-well culture plate, Sigma-Aldrich, Oakville, ON, Canada) and cell medium was replaced with low serum (1%FBS) media and culture was continued for additional 4 days. Spheroid monocultures were treated right away for 14 days. Spheroid co-cultures were treated for 21 days with media replaced every 4 days, where treatment was performed either from day7-10 or from day 7-21.

### 2.2. Therapeutic screening with Doxorubicin, o-Vanillin and RG-7112

2D: Doxorubicin hydrochloride (Sigma-Aldrich, #44583) solutions were prepared in PBS to obtain stock solutions. Working solutions were then diluted using low serum DMEM serving as the vehicle control and added to chambers at final concentrations of 0, 0.1, 0.25, and 0.5 µM Doxorubicin. 3D mono-/co-culture: Doxorubicin was used at a concentration of 0.25µM in monoculture and 0.5µM in co-culture. O-Vanillin solutions were prepared in PBS at stock concentration 0.02M to achieve a working solution of 100µM in full serum DMEM, and RG-7112 was prepared in PBS as a stock solution of 5mM, which was then diluted to a working concentration of 5µM in full serum media. Spheroids were treated for 14 days with one of the following: no drugs, Doxorubicin, or a combination of Doxorubicin, o-Vanillin, and RG-7112, all in low-serum media. In 3D co-culture experiments, drugs were diluted the same way. For the Doxorubicin induction experiment, concentrations of 0, 0.5, and 1.0µM Doxorubicin were used.

In the combination treatment experiment, a concentration of 0.5µM Doxorubicin was applied and the co-cultures were treated with either no drugs, Doxorubicin, Doxorubicin + o-Vanillin, Doxorubicin + RG-7112, or Doxorubicin + o-Vanillin + RG-7112, all in low-serum media.

### 2.3. Metabolic Activity Measurements

Metabolic activity was assessed with AlamarBlue assay as previously described [19] and as per the manufacturer’s instructions (Invitrogen, #DAL1100). Briefly, at the required time point, media is removed and low serum media with 10% AlamarBlue is added to each well and incubated at 37°C for 6 hours. Fluorescence exposure is set to Ex560nm/Em590nm and measured using a spectrophotometer (Tecan Infinite T200, Männedorf, Switzerland) using the Magellan software. All metabolic activity is shown as a percentage as compared to the experiments control. AlamarBlue assays were performed for the mono-culture spheroids on days 7 and 14 of treatment and for co-cultures after 3 and 14 days of drug treatment.

### 2.4. Immunohistological analysis

To determine *in vitro* efficacy of drug combinations on spheroids in mono- and co-culture experiments, we prepared the spheroids for sectioning and staining. 3D spheroid mono- and co-cultures were washed with PBS, and then fixed with 4% paraformaldehyde (Sigma-Aldrich, Oakville, ON, Canada), cryoprotected in 10-30% sucrose, and embedded in Tissue-Plus™ optimal cutting temperature compound (OCT) (Fisher Scientific, Canada). They were then flash-frozen in -80°C for cryopreservation. Sections were made at 16-µm-thick slices and mounted on Fisherbrand Superfrost™ Plus slides (Fisher Scientific, Canada) then placed in -20°C. Cryosectioning was performed on a Leica CM1950 Cryostat (Leica Microsystems, Richmond Hill, ON, Canada). Slides were placed on a 50°C heater for 30 minutes and then washed with PBS mixture with 0.1% Triton X-100 (Sigma-Aldrich, Canada) and 1% Tween (Sigma-Aldrich, Canada). Slides were then saturated in a blocking buffer made with a PBS-Tween-Triton mixture, containing 1% bovine serum albumin (Sigma-Aldrich, Canada) and 1% goat serum (depending on the host of the secondary antibody) for 1 hour at room temperature. For 2D, immunopositivity was done using a fluorescence antibody p21 (ab220206, Abcam, Cambridge, MA, USA) and then the slides were mounted with coverslips using Fluoroshield™ with DAPI (F6057-20ML, Sigma-Aldrich, St. Louis, MO, USA) with the same steps as mentioned above removing the step of exposure to heat. 3D cultures slides were incubated at 4°C overnight with the fluorescence antibody p21 (ab220206, Abcam, Cambridge, MA, USA), for senescence quantification and visualized using a fluorescent secondary antibody Alexa Fluor™ 555 anti-mouse (Invitrogen, ThermoFisher Scientific Waltham, MA, USA) and were then mounted with coverslips using Fluoroshield™ with DAPI (F6057-20ML, Sigma-Aldrich, St. Louis, MO, USA). For proliferation analysis, samples were incubated with primary Ki-67 rabbit mAb (ABclonal Technology, Woburn, MA, USA) at 4°C overnight and then Alexa Fluor™ 555 anti-rabbit (Invitrogen, ThermoFisher Scientific Waltham, MA, USA) conjugated secondary antibody for 2 hours at room temperature. Images were captured using an Invitrogen™ EVOS™ M5000 Imaging System (Invitrogen, #AMF5000) for both brightfield and fluorescent photomicrographs.

### 2.5. Tumor spheroid image analysis

Area of spheroids was measured using Fiji ImageJ version 1.0 (NIH, Maryland, USA) using ROI manager spheroid drawn area to measure and track spheroid growth over time. In the co-culture experiments, we applied a TopHat Fast Fourier Transform (FFT) filter (2.0 pixels) then inverse the FFT of the model image and measured fluorescence intensity and area under the curve from their corresponding 3D surface plot and ROI manager multi plot. Fluorescence intensity and area were measured at 3 days and 14 days post treatment.

### 2.6. Statistical Analysis

Data was analyzed using GraphPad Prism version 10 (GraphPad, La Jolla, CA, USA). Paired or unpaired t-tests were used when comparing two groups, and multiple pairwise comparisons (one-or two-way ANOVA) were run to evaluate multiple groups. All experiments were run four-five independent times with different osteoblast donors for per ‘*n*’ done in duplicate or triplicate. Statistical analysis was done with the *p*-value set to < 0.05 presenting data as +/-SEM.

## 3. Results

### 3.1. Senescence induction in MDA-MB-231 and primary osteoblast cells using Doxorubicin in 2D culture

To assess the impact of Doxorubicin (concentrations at or below the IC50) toward inducing senescence without inducing apoptosis, we first treated MDA-MB-231 cells in 2D culture with a dose-range of Doxorubicin (0.1-0.5µM) for 72h followed by immunostaining of p21 (an alternate marker for senescence since p16^INNK4a^ is deleted in these cells). MDA-MB-231 treatment with 0, 0.1, 0.25, and 0.5µM Doxorubicin resulted in 1.66 ± 0.5%, 5.90 ± 1.08 %, and 23.66 ± 7.79 %, p21 positive senescent cells, respectively, as compared to 1.51 ± 1.34 % in the control group. The results were statistically significant, as indicated in Figure 1. Treatment of primary human spine osteoblasts with the same concentrations of Doxorubicin in 2D culture for 72h also significantly increased p21 positivity of 14.77 ± 2.65 %, 24.31 ± 4.5%, and 47.95 ± 10.65%, respectively, as compared to 11.52 ± 3.25%, in vehicle treated controls.

**Figure 1.**
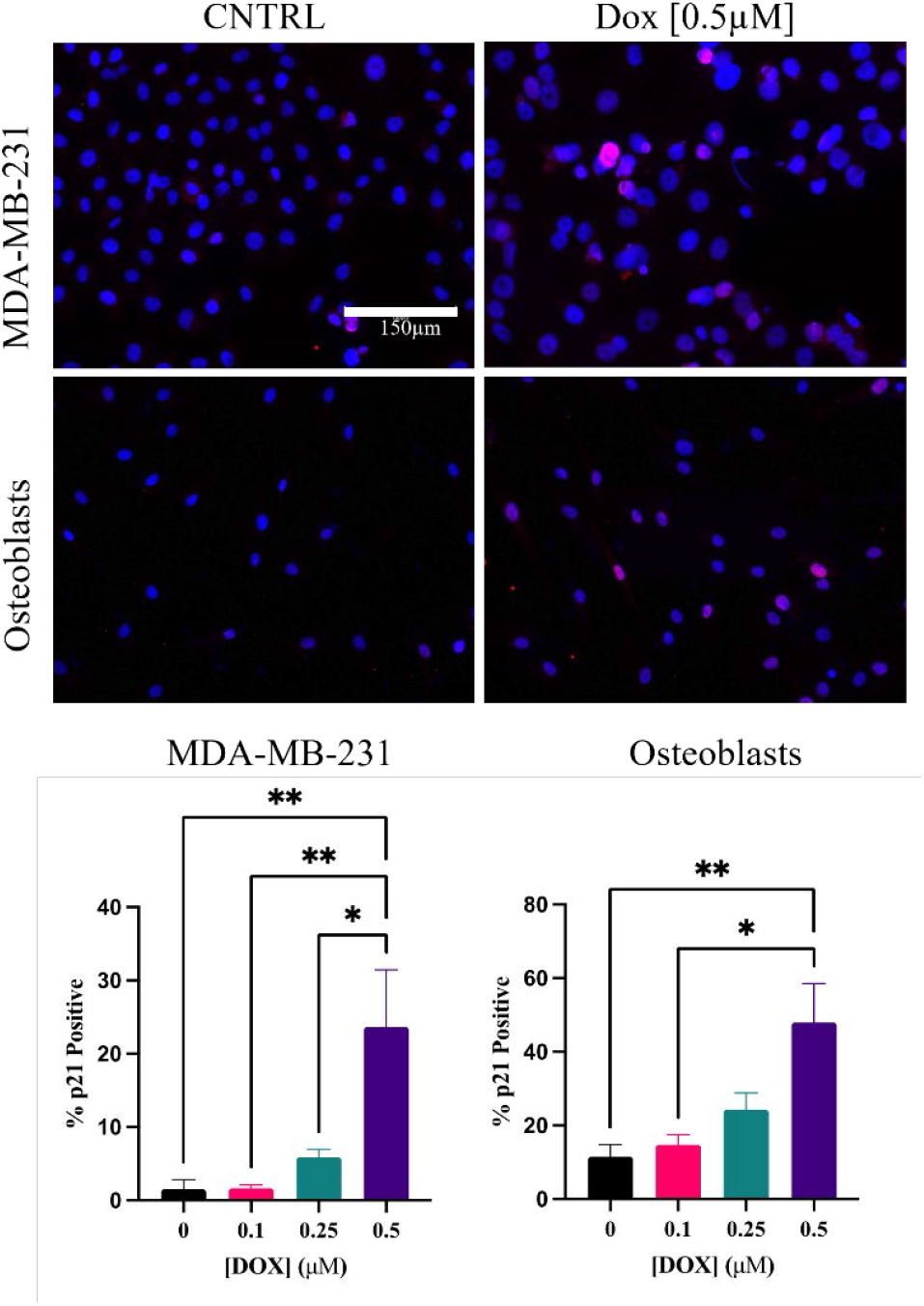
MDA-MB-231 and primary spine osteoblasts. Senescent activity evaluated by p21 immunofluorescence staining in 2D. Treated with 0-0.5µM Doxorubicin for 72h. Representative photomicrographs of p21 (red nuclear staining) expression in MDA-MB-231, and osteoblasts. DAPI staining with Fluoroshield was used for nucleus localization. Bottom graphs indicate quantification positive p21 stained MDA-MB-231 and primary osteoblast cells in 2D. n = 5, Mean +/-SEM, ^*^ = p < 0.05, ^**^ = p < 0.005, ^****^ = *p* < 0.0001, One-way ANOVA.

### 3.2. Senolytics combined with Doxorubicin significantly inhibit tumor spheroid growth and metabolic activity

To mimic the tumor microenvironment more closely, we assessed Doxorubicin-induced senescence in triple negative breast cancer cell spheroids. We further assessed the impact of senolytics on blocking Doxorubicin-induced senescence over a 14-day treatment period using a combination of 0.25 µM Doxorubicin, 100µM o-Vanillin and 5µM RG-7112. Fluorescence imaging of the GFP-expressing MDA-MB-231 cells was performed every 3 days (data not shown) and by day 14, combination of Doxorubicin and senolytics clearly showed that tumor sphere size was markedly less than Doxorubicin alone or vehicle treated samples (Fig 2A). Quantification of tumor sphere size revealed that spheroids treated with Doxorubicin combined with o-Vanillin and RG-7112 were significantly smaller in area compared to the vehicle control or Doxorubicin alone p < 0.0001 (Fig 2 B-C) 1.11 ± 0.029 fold, and the combination at 0.54 ± 0.04 fold as a ratio of control ^****^ = p < 0.0001. Metabolic activity assay revealed a significant reduction in viability in combination treatment samples of 0.41 ± 0.13 fold and 0.57 ± 0.10 fold for Doxorubicin alone compared to vehicle treated samples (Figure 2 D, p = 0.04). Taken together, these data indicate that combining senolytics o-Vanillin and RG-7112 increase the therapeutic efficacy of Doxorubicin evidenced through inhibiting tumor sphere growth and disrupting metabolic activity.

**Figure 2.**
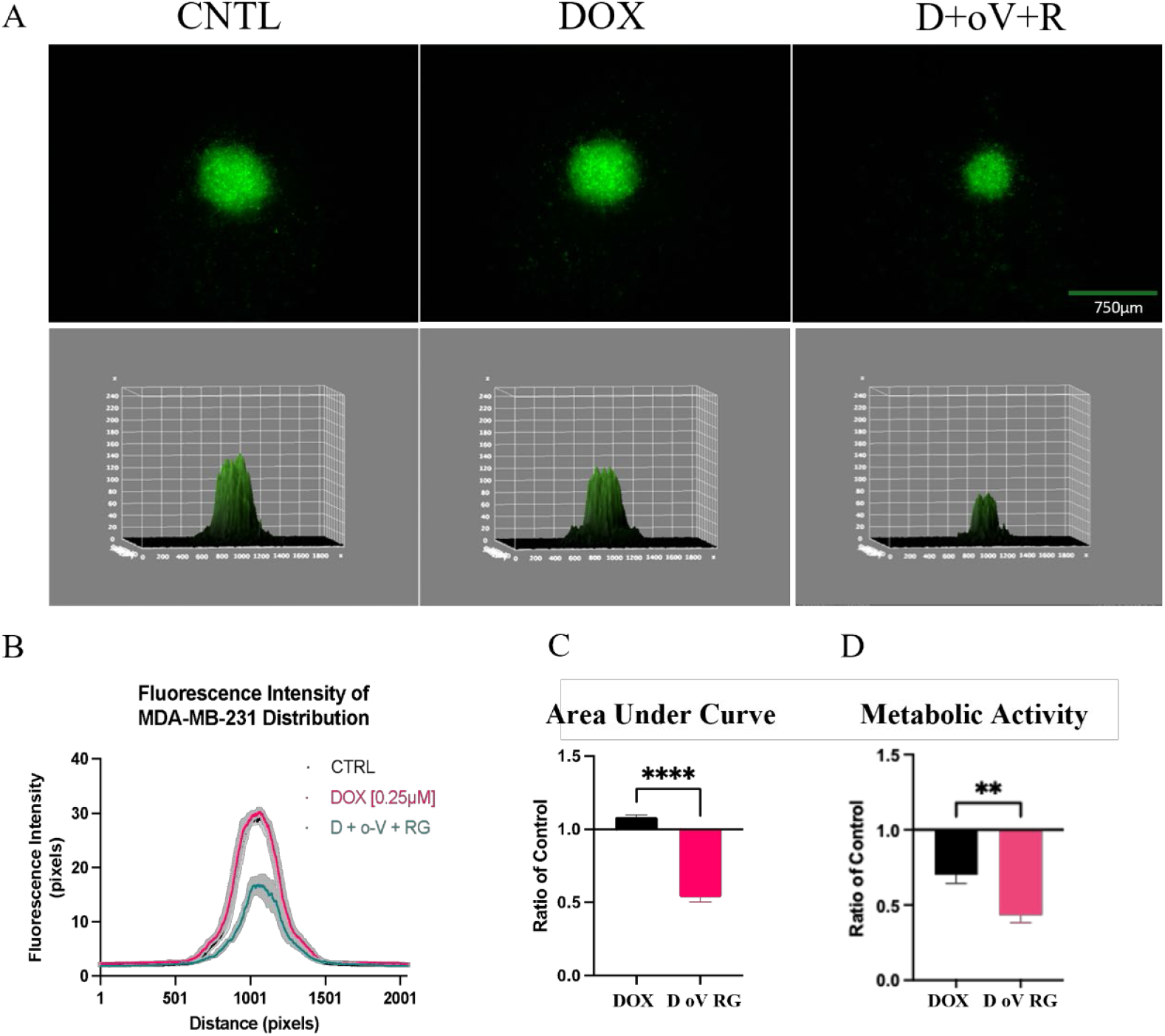
MDA-MB-231 spheroids treated 14 days with standard media as control, [0.25µM] Doxorubicin, or in combination with [100µM] o-Vanillin, and [5µM] RG-7112. **A)** Representative photomicrographs of GFP-tagged spheroids on day 14 of treatment with complimentary 3D surface and ROI multi plots. Quantification of **B)** Fluorescence intensity mean as a measure from a 3D surface plot **C)** Average area under the curve comparing Doxorubicin alone and combination treatment on day 14 to vehicle control n=4 **D)** Metabolic activity comparing Doxorubicin alone and combination treatment on day 14 to vehicle control n=3, Mean +/-SEM. ^**^ = p = 0.04, ^****^ = *p* < 0.0001, unpaired t test.

### 3.3. Doxorubicn induced senescence in a model of the bone metastatic niche

To evaluate senescence induction in a bone-like tumor microenvironment we used a co-culture 3D model with GFP tagged MDA-MB-231 cells as the center spheroid, surrounded by primary spine osteoblasts in a type I collagen matrix. A preliminary dose-response evaluation indicated that higher concentrations of Doxorubicin were necessary to induce the same level of senescence as in 2D culture experiments (data not shown). We tested a range, and found in 3D culture the 0.25µM was insufficient to induce senescence and 0.5µM was (data not shown). Therefore, we increased the dose to 0.5µM Doxorubicin. This more complex 3D co-culture was then treated with either Doxorubicin alone or in combination with the senolytics to evaluate senescence blockade and additive efficacy. **Figure 3** shows immunohistochemistry staining of p21 positive senescent cells of each cell type in the co-culture. Treatment with Doxorubicin significantly induced senescence in both cell types. For MDA-MB-231 cells, Doxorubicin increased p21 positive cells to 8.31 ± 1.86% compared to 1.17 ± 0.41% p21 positive in controls. Addition of the two senolytics significantly inhibited p21, reducing it to 0.98 ± 0.4% p21 positive cells. For the primary osteoblasts (cell lacking GFP expression, observed in the peripheral collagen gel), Doxorubicin increased p21 positive cells to 12.95 ± 1.64% compared to 2.53 ± 0.66% p21 positive in controls. Addition of the two senolytics significantly reduced p21 to 3.91 ± 1.72% positive. This data shows that in a more complex 3D tumor-like microenvironment, Doxorubicin induced senescence in both the tumor and stromal compartments. Moreover, the addition of the two senolytics brought the senescence levels back to baseline in both compartments.

**Figure 3.**
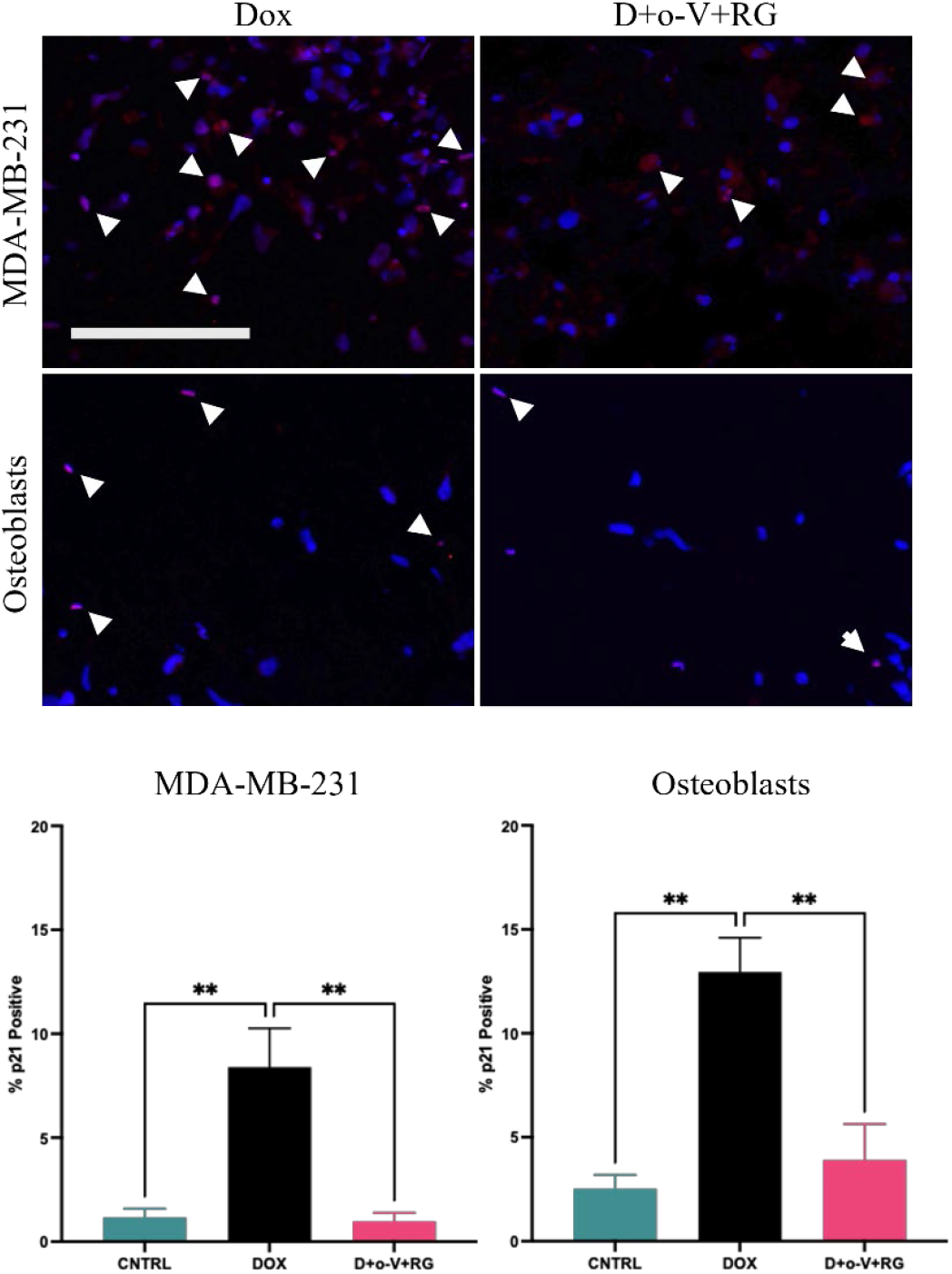
3D culture of MDA-MB-231 spheres surrounded by primary osteoblasts in a collagen gel. Samples were treated with Doxorubicin [0.5 µM] alone or in combination with o-Vanillin [100µM] and RG-7112 [5µM]. The representative micrographs show p21 positive MDA-MB231 and primary osteoblasts indicated by arrow heads. Quantification of p21 positive staining after 14 days of treatment is shown in the corresponding bar graphs. Mean +/-SEM. ^**^ = p < 0.005, one-way ANOVA, n=4.

### 3.4. Addition of senolytics disrupts tumor sphere growth in a bone metastatic model

To evaluate the impact of senolytics on tumor outgrowth in the bone metastatic niche, the co-culture model was treated with Doxorubicin alone or in combination with the two senolytics over 2 weeks. Figure 4A shows representative images of GFP-expressing breast cancer spheres over the course of treatment, with their corresponding fluorescence intensity ROI multi-plots. After 3 days of treatment with Doxorubicin alone or with the combined treatment, tumor sphere sizes were similar based on intensity plots (Figure B), however, spheres were observably smaller for samples treated with Doxorubicin and more so for combined treatment after 14 days. Quantification of sphere area at 14 days indicated that Doxorubicin-treated samples were significantly smaller (0.70 ± 0.06) compared to controls. The combined treatment had an added effect and further significantly reduced the size (p < 0.05), 0.43 ± 0.05) compared to the control. This indicated that combined treatment was about 1.65 fold smaller sphere than Doxorubicin alone. After 3 days of treatment, a non-significant reduction in viability in Doxorubicin and combination treated samples compared to vehicle treated controls (Figure 4 D). After 14 days of treatment, there was a significant reduction in viability, in Doxorubicin treated samples (0.60 ± 0.05) and in combination treated samples (0.09 ± 0.06) compared to vehicle treated controls (Figure 4 C, p < 0.001), which represented approximately a 6.7-fold decrease in viability of combined treatment compared to Doxorubicin alone. This indicates that combining senolytics o-Vanillin and RG-7112 with Doxorubicin significantly inhibits triple negative breast cancer spheroid growth, cell outgrowth and metabolic activity compared to Doxorubicin alone and untreated controls.

**Figure 4.**
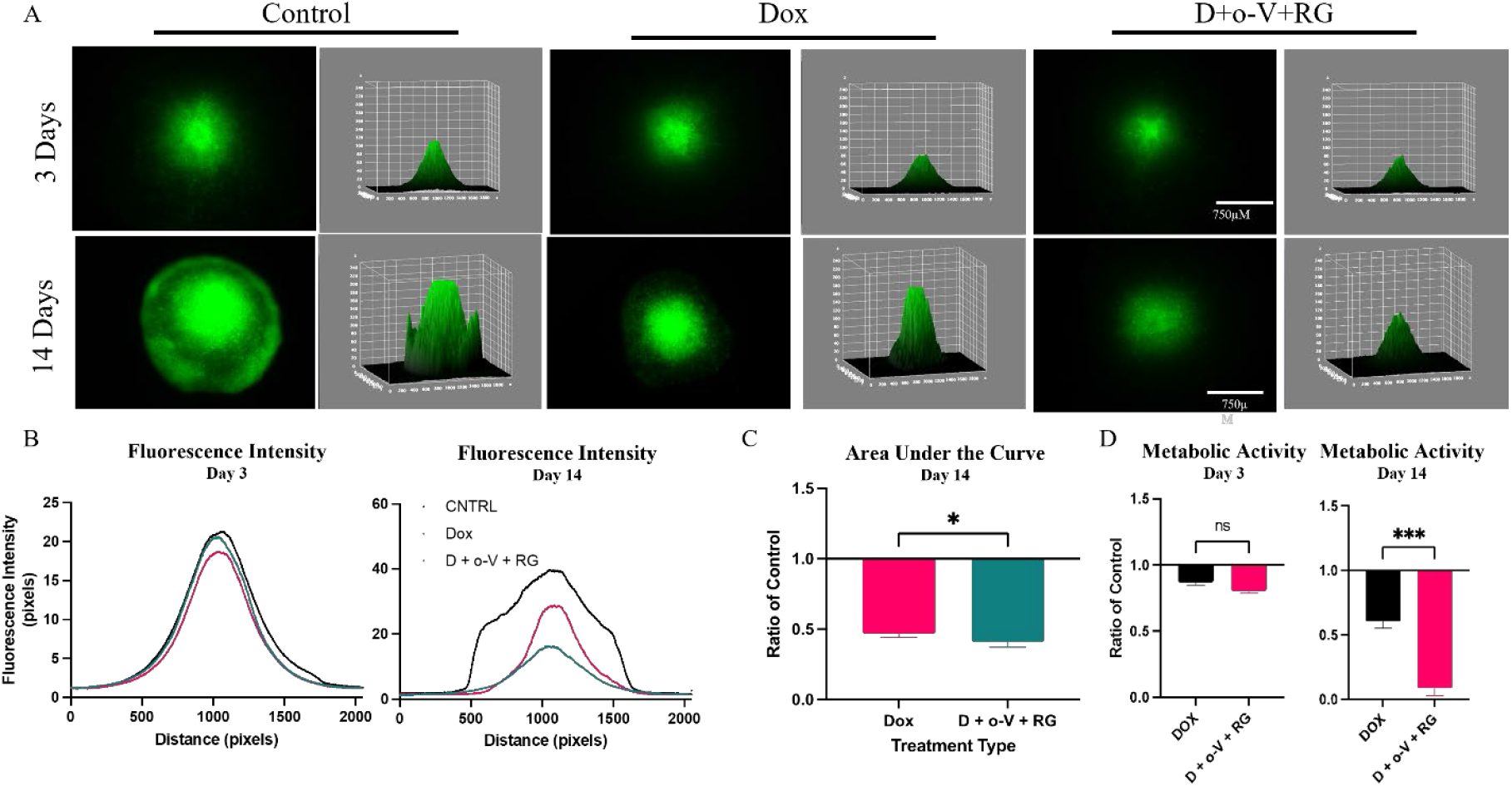
Tumor growth in a bone metastatic microenvironment treated with Doxorubicin alone or in combination with o-Vanillin and RG-7112. **A)** Representative photomicrographs of GFP-tagged spheroids with complimentary 3D surface and ROI multi plots at 3 and 14 days of treatment. B) Tumor spheroid fluorescence intensity at 3 and 14 days of treatment. C) Quantification of area under the curve as the ratio of control. D) Metabolic activity of Doxorubicin alone and the combination treatments as a ratio of control at days 3 and 14 post-treatment. N = 4, Mean +/-SEM.^***^ = p < 0.001, one-way ANOVA, unpaired t-test.

**Figure 5.**
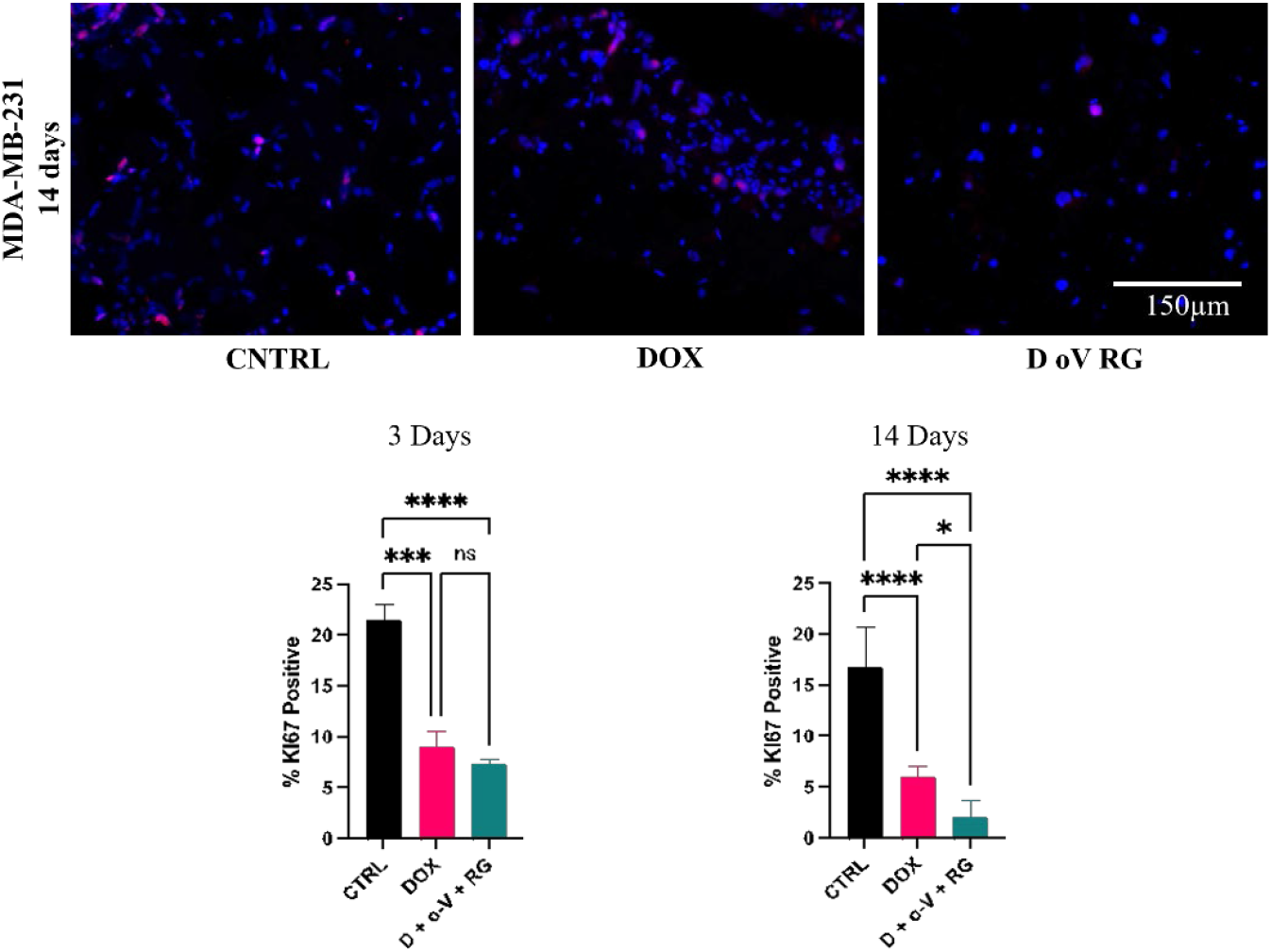
Assessment of tumor cell proliferation in a bone metastatic model following treatment with Doxorubicin or combination treatment. Immunohistochemistry staining for Ki67 positive cells was performed at 3 and 14 days post-treatment. A) The total number of cells (blue) and Ki67 positive cells (red) for each condition on day 3 of treatment. **B)** % Ki-67 positive on day 3. **C) D)** % Ki-67 positive on day 14. n = 4, Mean +/-SEM. ^*^ = *p* < 0.05, ^****^ = p < 0.0001, 2way ANOVA.

### 3.5. Combining senoltyics with Doxorubicin reduces proliferation of tumor cells a bone metastatic model

Cellular proliferative activity was assessed immunohistochemically, staining for the marker Ki-67. After 3 days, Doxorubicin-treated samples displayed 9.41 ± 1.51 % Ki67 positive cells, combination-treated samples displayed 7.36 ± 0.46 % Ki67 positive cells compared to 21.54 ± 1.53 % Ki67 positive cells in the vehicle-treated control. While both treatments were significantly less than the controls, combined treatment was not significantly different from Doxorubicin alone. After 14 days, Doxorubicin-treated samples displayed 6.03 ± 0.82 % Ki67 positive cells, combination-treated samples displayed 2.12 ± 1.18 % Ki67 positive cells compared to 16.89 ± 2.71 % Ki67 positive cells in the vehicle-treated control. Both treatment groups had significantly less positive Ki67 than vehicle-treated control, and the combined treatment had a significant additive effect over Doxorubicin alone (p < 0.05). Taken together, the data suggest that adding senolytics can potentiate the anti-proliferative effect of Doxorubicin against triple-negative breast cancer cell growth in a bone-like microenvironment.

## 4. Discussion and Conclusion

In the present study, we hypothesized that selective removal of Doxorubicin-induced senescent tumor and stromal cells would increase overall Doxorubicin efficacy. Originally, senescence was deemed non-harmful in relation to cancer as the cells are non-proliferative. However, with time it was understood that in accumulation these cells adopt the SASP phenotype and support an environment of proinflammatory cytokines and chemokines allowing for further growth of the primary cancer. Low doses of Doxorubicin are used to reduce side-effects in patients; however, they are known to induce cellular senescence. As such, we evaluated if clinically relevant doses of Doxorubicin would induce senescence in both cancer and stromal cells in standard 2D and in a 3D bone-like microenvironment. Two different senolytics were then tested for their ability to reduce the senescent cell burden and positively impact the efficacy of Doxorubicin. We found that clinically translatable Doxorubicin doses induced senescence in both triple-negative human breast cancer cells and primary human spine osteoblasts, in both 2D culture and 3D spheroid culture. Furthermore, we show that a combination of the senotherapies o-Vanillin and RG-7112 in combination with Doxorubicin significantly reduced the number of senescent cells, reduced metabolic activity, proliferation and tumor sphere growth when cultured alone or in co-culture with primary spine osteoblasts. This was correlated with significantly reduced proliferation in combination-treated samples. Showing anti-cancer additive effects using clinically relevant concentrations of senolytics. These results suggest that removing both senescent stromal and tumor cells can significantly improve the effects of Doxorubicin treatment. Therefore, combining senolytic therapies may have a large impact on improved health outcomes for patients undergoing this type of cancer treatment.

Chemotherapeutics, such as Doxorubicin, are a standard treatment for both primary and metastatic breast cancers. Emerging evidence suggests that current effective clinical doses of Doxorubicin (and others) can induce cellular senescence[32, 33] in primary breast cancer, in bone/spine metastatic disease secondary to breast cancer, and within the stroma/microenvironment. [34]. SASP factors produced by these cells are thought to disrupt immune function and drive drug resistance and tumor progression. To assess whether o-Vanillin and RG-7112 could target and remove Doxorubicin-induced senescent triple negative breast cancer cells and/or primary osteoblasts, we evaluated if clinically relevant Doxorubicin concentrations could reproducibly induce senescence. Since blood concentrations of patients treated with Doxorubicin are typically between 0.025 – 0.25 µM (as opposed to many *in vitro* studies using 1-5 µM) [35], we used these lower clinically relevant doses to mimic *in vivo* dosing and induce senescence in our study. Our data confirmed that Doxorubicin doses within this range significantly increased triple negative breast cancer and primary human osteoblast senescence, as indicated by significant increases in p21 positive staining in both 2D and 3D cultures. This was the first time to our knowledge that Doxorubicin was shown to induce human primary osteoblast senescence. Similar studies were performed in human chondrocytes [36] or animal bones [37], where in both studies, Doxorubicin was used as the inducer of senescence with similar dosing (0.1 µM – 1 µM). Our findings are able to help establish a perhaps more physiologically representative *in vitro* model of human triple negative breast cancer cells in a human bone-like microenvironment.

As mentioned, clinical doses of Doxorubicin are designed to reduce off-target effects, yet this type of exposure has also been shown to cause therapy-induced senescence [32, 33] in primary breast cancer and in bone/spine metastatic disease secondary to breast cancer[34]. While induction of senescence was originally thought to benefit the patient by inhibiting tumor growth and allowing immune cells to remove the target tumor cells. It is now recognized that senescence in the tumor and surrounding stroma paradoxically promote tumor progression, likely attributed to the secreted SASP [38, 39]. For instance, targeting senescent cells in the stroma could improve fracture healing and tissue homeostasis by reducing SASP [40]. Removal of senescent cells (using Quercetin) in the bone *tumor microenvironment* may therefore improve bone health and block tumor progression [41, 42] through reduced inflammatory cytokines. While cytokine and SASP involvement was not assessed in the present study, it can be speculated that removal of senescent osteoblasts and breast cancer cells would reduce SASP factors thereby improving Doxorubicin efficacy. Taken together with our findings in the present study, combining senolytics with clinically relevant chemotherapy doses may help improve the treatment of bone related malignancies and promote bone health.

RG-7112 is a synthetic peptide derived from a previous synthetic senolytic with anti-cancer activity, nutlin-3a [26]. It is a p53-MDM2 inhibitor that restores p53 activity, was originally developed for anti-leukemia treatment and tested in phase-1 clinical trials [28]. Although RG-7112 failed to pass phase 2 trials due to adverse side effects [43], lower and safe doses of RG-7112 combined with o-Vanillin can effectively remove senescent cells from other human and animal musculoskeletal tissues [29, 44, 45] and reduce pain and restore disc height in an animal model of back pain [45]. The 100µM o-Vanillin used in combination with 5µM RG-7112 in the present study showed high promise for blocking triple negative breast cancer growth and migration in both 3D spheroids and a 3D co-culture system. Interestingly, a recent study showed that RG-7112 at 6µM could not effectively reduce sphere size or viability of both triple negative and hormone responsive tumor spheres or patient derived triple negative tumor spheres[46]. However, it is important to note that treatment times went for a maximum of 72 hours, and no additional treatment of chemotherapeutics or additional senolytics were given. Our preliminary findings also suggested that RG-7112 could only moderately disrupt the tumor cell growth or metabolic activity on its own, however, significantly disrupt tumor cell growth when in combination with o-Vanillin. O-Vanillin is a metabolite of Curcuma, and it is known to have anti-inflammatory, senomorphic and anti-cancer properties [29, 47, 48]. A recent study from 2022 indicated that a derivative of o-Vanillin displayed an IC50 of around 40-50 µM against human osteosarcoma, triple negative and hormone responsive breast cancer cell lines in 2D culture. This was also linked with increased measure reactive oxygen and increased apoptosis[48]. Those cells were, however, not induced with any chemotherapy, and the basal senescence level was not assessed. Our preliminary work indicated that o-Vanillin on its own (100 µM) had no impact on Doxorubicin-induced triple negative breast cancer cell growth or viability in mono-culture spheres or in 3D co-culture. In terms of pre-clinical animal studies, co-administration of senolytics with chemotherapeutics has been shown to increase longevity and reduce tumor burden by removing senescent cells. Navitoclax, a natural anti-inflammatory senolytic/senomorphic, combined with cisplatin [49] at low doses was confirmed to kill senescent cells with no adverse effect on normal cells, all while slowing the growth of tumor compared to control treatments of cisplatin or Navitoclax alone. Therefore, we can envision future work testing o-Vanillin and RG-7112 combined with Doxorubicin *in vivo* for increased efficacy against primary breast cancer and metastatic bone disease xenografts.

In this study we set out to assess the impact of combining senolytics with Doxorubicin to enhance efficacy against triple-negative breast cancer growth in a bone metastatic niche. Previous work from our group demonstrated that a combination of o-Vanillin and RG-7112 could effectively remove senescent cells, reduce inflammatory SASP, block painful tissue degeneration and improve tissue homeostasis *in vivo* and *in vitro* for musculoskeletal degenerative disease[22, 50]. Considering chemotherapies are known to induce senescence[38] and senescent cells within the tumor and surrounding stroma are known to contribute to therapy resistance and cancer progression [20], we hypothesized that combining o-Vanillin and RG-7112 with a standard chemotherapeutic Doxorubicin would enhance treatment efficacy. We showed that Doxorubicin indeed induces senescence in both triple negative breast cancer cells as well as primary human spine osteoblasts. Further, we showed that adding the two senolytics could reverse senescence and block tumor sphere growth and metabolic activity in 3D spheroid and co-cultures of spheroids in a bone-like microenvironment. This was correlated with a significant reduction in cell proliferation. Our results are compelling; however, further experiments using additional subtypes of breast cancer are required since the genetic profiles of hormone responsive or HER2+ subtypes are different to those of triple-negative breast cancers. It would also be beneficial to focus on patient derived bone metastases secondary to breast cancer to gain a more clinically relevant insight into this combined therapy. While our previous work has demonstrated that o-Vanillin and RG-7112 are tolerated well *in vivo* [50], future work should also focus on testing these drugs in combination with systemic Doxorubicin in both primary and secondary breast cancer *in vivo* models. Taken together, our results indicate strong promise for combining senolytics with systemic chemotherapy to reduce therapy-induced senescence and further inhibit tumor progression and metastasis.

